# Are plants in sown flower strips suitable for communities of wild bees? Pollination network approach in conservation biology

**DOI:** 10.1101/2021.05.29.446282

**Authors:** Jiří Hadrava, Anna Talašová, Jakub Straka, Daniel Benda, Jan Kazda, Jan Klečka

## Abstract

1. Drastic reductions of insect diversity and abundance are observed in the highly fragmented agricultural landscapes of central Europe. Declines of pollinators may have detrimental effects on the reproduction of wild insect-pollinated plants as well as the yield of crops. In order to mitigate such impacts, sown flower strips on arable land within Agri-Environment Climate Schemes (AECS) are supported across EU countries. However, it is not clear whether sown flower strips provide equivalent benefits to wild flower-visiting insects as semi-natural habitats.
2. Here, we apply plant-pollinator network approach to evaluate the function of sown flower strips for the communities of wild bees. We compared the structural characteristics and the robustness of plant-pollinator networks in sown flower strips and nearby semi-natural habitats. We also quantified the importance of individual plant species for bees based on simulations of plant-pollinator extinction cascades.
3. We found that assemblages of plants and pollinators were less diverse in sown flower strips than in semi-natural habitats, more generalized, and more nested. However, we did not find any significant differences in network robustness to plant-pollinator coextinctions. Further, simulations revealed a large variation in the functional importance among plant species from both habitats.
4. We conclude that although the analysis of network robustness suggested that plants in the sown flower strips and semi-natural habitats were functionally equivalent, this masked important differences among the two habitats. From the conservation point of view, semi-natural habitats were superior in supporting a more diverse community of solitary bees and bumblebees.

## Introduction

Habitat loss caused by intensive agricultural practices has been identified as the main driver of mass declines of the abundance and diversity of insects (Hallmann et al. 2017, Sánchez-Bayo & Wyckhuys 2019, Hallmann et al. 2021), including pollinators such as wild bees (Ollerton et al. 2014), in many regions over the last several decades. Homogenization of agricultural landscapes and consequent fragmentation of natural habitats is also one of the biggest threats to insect biodiversity in central Europe (e.g. Ekroos et al. 2010, Šlancarová et al. 2014). Among other ominous issues, insect declines may negatively affect pollination of wild plants (Jennersten 1988, Ollerton et al. 2011) as well as cultivated crops resulting in reduction of agricultural yields (Aizen et al. 2009, Garibaldi et al. 2013). Wild bees, such as solitary bees or bumblebees, are an important group of pollinators and their populations are strongly endangered in present-day European agricultural landscape by a combination of the lack of floral resources and nesting habitats and exposure to pesticides (Biesmeijer et al. 2006).

Reversing the decline of bees and other pollinators in agricultural habitats requires changes in agricultural practices (Klein et al. 2007, Potts et al. 2016). These include a shift to organic farming, which eliminates the use of pesticides, plus adding measures to increase the availability of floral resources, such as establishing sown flower strips at the field margins (Haaland et al. 2011, Scheper et al. 2013, Marja et al. 2018). Farmers across Europe have been financially motivated to sow flower strips on arable land within Agri-Environment Schemes (AES) since 2001 (Haaland et al. 2011), renamed as Agri-Environment Climate Schemes (AECS) since 2015. The design of sown flower strips within AECS varies among EU member states. They range from annual stands sown with economically accessible crops attractive to pollinators, e.g. *Phacelia tanacetifolia, Trifolium* spp., *Fagopyrum esculentum* (Tschumi et al. 2015, Wood et al. 2015), to perennial mixes similar to former European unimproved meadows containing many species of wild plants (Natagriwal 2018).

In the Czech Republic, AECS ‘nectar-rich flower strips’ (‘nektarodárné biopásy’) were introduced in 2015. They are sown in the spring with a uniform seed mix containing annual flowering crops as well as perennial species dominated by *Trifolium* spp. They should be maintained for two or three years and then reestablished at the same place or at a different location within the farm for the next period of two or three years. Once a year they should be cut any time between July 1 and September 15 (Ministry of Agriculture of the Czech Republic 2018). As no split management is required, the flowering period is not continuous throughout the season (Šrámková 2014). In the early summer of the seeding year, when the sown flower strips are at the peak bloom, they are highly attractive mainly for honeybees, bumblebees and hoverflies, and occasionally for solitary bees, which all feed especially on *Phacelia tanacetifolia*. Later in the season, the dominant flowering plant is *Melilotus albus*, visited mostly by honeybees. In the second year, *Trifolium* spp. prevails in the flower strips, which makes them attractive for honeybees and bumblebees (Šrámková 2014).

Numerous studies have shown that the presence of sown flower strips along field margins benefits the diversity and abundance of pollinators compared to farms without flower strips (Scheper et al. 2015, Buhk et al. 2018, Marja et al. 2018). The positive effect of flower strips is especially pronounced in landscapes with low availability of floral resources (Scheper et al. 2015). It is well established that the abundance of bees and other insects tends to be higher in the flower strips compared to crops (Haaland et al. 2011). However, it is less clear how the flower strips compare to semi-natural habitats in the surrounding landscape, even though there is evidence that the abundance and species richness of pollinators may be similar or higher in flower strips compared to extensive grasslands (Scheper et al. 2013). Species richness and composition of the plants included in the seed mixture also plays an important role in the attractiveness of flower strips for pollinators (Carreck & Williams 2002). While the diversity of pollinators in sown flower strips generally increases with plant diversity (Scheper et al. 2013), some pollinators show an increase in abundance in the presence of specific plant species, e.g. bumblebees are known to benefit from the inclusion of Fabaceae, such as clover (*Trifolium* spp.) (Carvell et al. 2006). On the other hand, a previous study by Wood et al. (2017) suggested that sown flower strips may be of limited value to solitary bees; they showed that only a minority of solitary bee species visited plants in the flower strips compared to seminatural habitats.

Until now, studies have focused on the abundance and diversity of pollinators in sown flower strips, but not much attention has been paid to the structure of entire plant-pollinator networks in the sown flower strips (Ouvrard et al. 2018) and to the comparison to semi-natural grasslands. Pollinators do not feed on plant species at random, but they exhibit a complex foraging behaviour leading to specialized or generalized interactions between plants and their pollinators (Waser et al. 1996), driven to a large extent by the selectivity of different pollinator species for various floral traits of the plants (e.g. Junker et al. 2013, Klecka et al. 2018b). Thus, the composition of flowering plants in the plant assemblage determines the suitability of the habitat for pollinators. However, also the structure of the relationships within the ecological network provides information which can be used to guide efforts at the conservation of biodiversity (Tylianakis et al. 2010). For example, Devoto et al. (2012) studied the assembly of pollination networks during the succession process and identified key plant species for the restoration of pollination networks in the model ecosystem of Scottish woodlands. In another study, Biella et al. (2017) identified which plant species are important for the conservation of grassland ecosystems in Northern Apennine mountains based on the analysis of modularity in pollination networks. Similar methods could be used to evaluate the effect of sown flower strips on pollinator communities.

An approach which has been hailed as a promising tool to obtain insights for conservation is based on evaluating the robustness of entire plant-pollinator networks to species extinctions (Memmott et al. 2004). In ecological networks, an extinction of one species may lead to coextinctions of other species which depend on it (Dunn et al. 2009). Memmott et al. (2004) developed the first model for simulations of coextinctions in plant-pollinator networks based on the assumption that a pollinator species becomes extinct when it loses all species of plants whose flowers it normally visits and a plant goes extinct when it loses all pollinators. Quantitative evaluation of the robustness of the network is possible based on the area under the simulated coextinction curve (Burgos et al. 2007). Further research of animal coextinctions driven by disappearance of plant species was then generalized for a wider range of animal groups (Pocock et al. 2012), for heterogeneous structure of habitats (Evans et al. 2013), and for temporally dynamic networks (Kaiser-Bunbury et al. 2010). The method has also been used for predictions of coextinctions driven by climate change (Memmott et al. 2007, Schleuning et al. 2016). Because these simulations of coextinctions are based only on the topology of the plant-pollinator mutualistic network, this model is called *topological coextinction model* (TCM) according to Vieira & Almeida-Neto (2015), who developed a different, so-called *stochastic coextinction model* (SCM). The SCM assumes that the secondary extinctions are stochastic and their probability increases proportionally with increasing dependency of individual species on the previously extinct species of the other trophic level.

Here, we are using both approaches. We agree with arguments given by Vieira & Almeida-Neto (2015) that unlike TCM, SCM brings biologically relevant assumptions such as higher importance of more frequent interactions and the ability to survive on alternative resources. However, we argue that TCM has also still some interpretative advantages over SCM: the SCM assumes that each species depends on its partners proportionally to the observed frequencies of interactions between them, so it does not take into account the flexibility of species (in our case pollinators) to shift their interactions after the extinction of some of their previous partners (visited plants). Such flexibility has been experimentally documented for example by Biella et al. (2019, 2020). On the other hand, TCM assumes the opposite extreme, i.e. that a species survives until any single partner of the species remains present. Despite their conceptual differences, both coextinction models received comparable empirical support based on field experiments in grasslands in central Europe (Biella et al. 2020). Moreover, the simulations can be used to quantify the functional importance of individual species in the network (Pocock et al. 2012).

In this study, we used analyses of the structure of plant-pollinator networks and simulations of coextinctions of plants and pollinators to compare the suitability of sown flower strips and semi-natural grasslands for aculeate Hymenoptera, a major group of pollinators. Specifically, we asked i) whether sown flower strips host a comparable diversity of Aculeata as nearby semi-natural grasslands, ii) whether the structure of the plant-pollinator networks and consequently their robustness to species extinctions systematically differ between the two habitats, and iii) whether individual plant species in the sown flower strips have higher or lower level of functional importance than plants in semi-natural grasslands based on simulations of coextinctions of plants and pollinators.

## Materials and methods

### Study sites

Plant-pollinator networks were studied at 7 sites with sown flower strips, established in the spring of 2016 using a mixture of seeds of 13 plant species (Table 1). Each site was situated in an ordinary agricultural landscape of central Bohemia, Czech Republic (K: 50.1187N, 14.2311E; M: 49.6029N, 14.2545E; P: 49.5478N, 14.3584E; R: 50.0878N, 14.2990E; S: 50.0086N, 14.8779E; U: 50.0360N 14.6192E; Z: 49.5496N, 14.9579E, see map in Fig. 1). At each site, two sampling plots were established: one in a sown flower strip and one in a semi-natural herbaceous habitat with occasional shrubs or trees (mostly uncultivated land between fields) with the highest diversity of dicotyledonous plant species in the diameter of 700-1000 m far from the flower strip. The sown flower strips were set up either at field margins adjacent to semi-natural habitats in the surroundings of the fields, or up to 60 m from the nearest semi-natural habitat. The strips varied in size; their length ranged from 100 to 400 meters and their width ranged from 3 to 12 meters. They also differed in mowing regimes; four of them were completely mown once at various times during the season and three of them were mown twice per year, but only a part of the strip was mown each time.

**Table 1.** The list of plant species used to establish the sown flower strips. The weight of seeds sown per hectare is provided.

| Family | Species | Seeds (kg/ha) |
| --- | --- | --- |
| Brassicaceae | *Sinapis alba* | 1.5 |
| Polygonaceae | *Fagopyrum esculentum* | 2.5 |
| Boraginaceae | *Phacelia tanacetifolia* | 1 |
| Fabaceae | *Trifolium pratense* | 4 |
| Fabaceae | *Trifolium hybridum* | 0.4 |
| Fabaceae | *Trifolium repens* | 0.4 |
| Fabaceae | *Melilotus alba* | 1 |
| Fabaceae | *Lotus corniculatus* | 0.2 |
| Fabaceae | *Onobrychis viciifolia* | 5 |
| Fabaceae | *Vicia sativa* | 5 |
| Apiaceae | *Carum carvi* | 2.5 |
| Asteraceae | *Achillea millefolium* | 0.01 |
| Malvaceae | *Malva sylvestris* | 0.05 |
| **Total** |  | **23.56** |

**Fig. 1.**
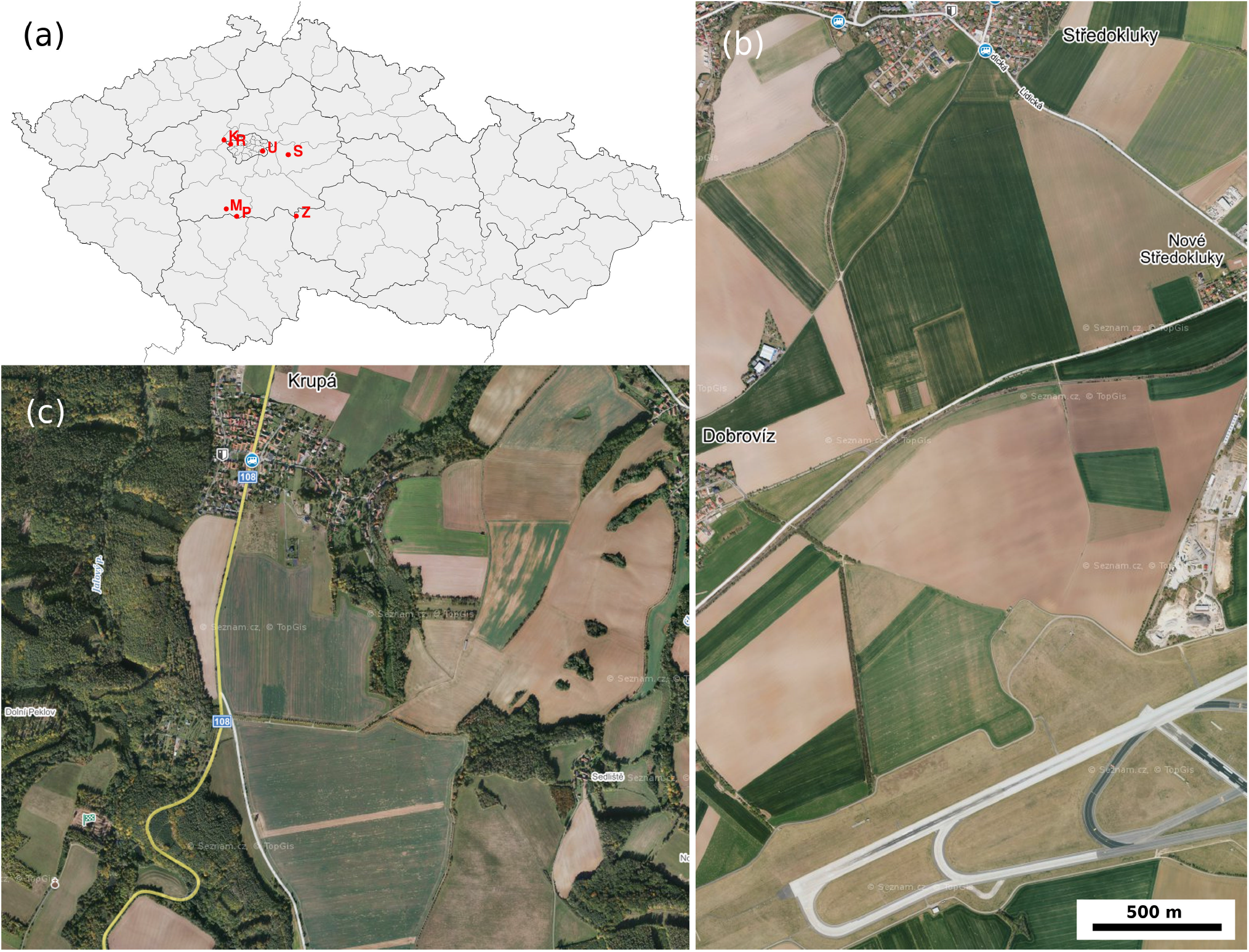
Location of the study sites (a) and aerial photos of two examples of the study sites: site K (b) and site S (c).

### Sampling protocol

Plant-flower visitor interactions were sampled in 2016, in three summer terms at monthly intervals from the end of June to the beginning of September. The method of standardized transect walks was used to detect plant-pollinator interactions. At each site, transect walks were done in a corridor of 100 m x 1 m between 9 A.M. and 4 P.M., during suitable weather conditions for the studied species: minimum of 18 °C, low wind, no rain, and dry vegetation. In order to have uniform collector bias throughout the study, all transect walks were done by one surveyor (AT). Honeybees and bumblebees were counted in the field and identified at the species level except for individuals of *Bombus terrestris* and *B. lucorum* that are indistinguishable in the field and thus, they were treated as single taxon. Additionally, based on records from yellow pan traps (data are not presented in this paper), we found that only 0.2 % of specimens from the *B. terrestris/lucorum* group belong to *B. lucorum* at the study sites. Other bee species that could not be identified in the field were collected with a sweep net for later identification, provided by JS and DB. All sampled material is stored in the National Museum in Prague.

### Network analyses

Data were analysed in software R, version 3.2.3. (R core team 2016). Pollination network analyses were done with package ‘bipartite’ (Dormann et al. 2008), mixed effect models were done with package ‘lme4’ (Bates et al. 2014) and diversity indices were calculated using package ‘vegetarian’ (Charney & Record 2012).

For each site, 3 plant-pollinator interaction networks were constructed based on observations pooled from the three sampling dates: i) the network of interactions observed in the sown flower strip only, ii) the network of interactions observed in the semi-natural habitat only, and iii) the overall network merging observations from both habitats.

Divided networks (i and ii) were used for the comparison of metrics of network structure between the sown flower strips and the semi-natural habitats. For each network, we counted the number of species of animals (A) and plants (P), calculated web asymmetry (Blüthgen et al. 2007), nestedness measured as weighted NODF (Almeida-Neto & Ulrich 2011), network specialization (Blüthgen et al. 2006), and the robustness (R) of the network to the secondary extinctions based on 10000 simulations of coextinctions under the conditions of the topological coextinction model (TCM) (Burgos et al. 2007) and the stochastic coextinction model (SCM) (Vieira & Almeida-Neto 2015). The differences of the metrics of network structure between networks from sown flower strips and networks from semi-natural habitats were tested with paired t-tests on original data or, in the case of A, P and wNODF that have positively skewed distribution, on log-transformed data. Moreover, we calculated the Shannon index of beta-diversity, separately for plant and bee data, to evaluate the variation of species composition in the two habitat types between sites (Jost 2007).

The cumulative networks covering the community from the whole site (iii, i.e. both habitats combined) were used for the evaluation of the importance of individual plant species for the whole local community of bees. For this purpose, we calculated several species-level indices for each plant species: normalised node degree (ND) (Dormann 2011), standardized specialization d’ (Blüthgen et al. 2006) and pollinator support index (PSI) (Dormann 2011), which is identical to species strength defined by Bascompte et al. (2006) as the sum of the dependence values of the pollinators on the plant. We compared the values of species-level indices of plants growing in the sown flower strips with plants growing in the semi-natural habitats using generalised linear models (GLMM) with Gaussian error distribution and site identity as a factor with random effect. The ND and PSI were log-transformed prior to the analysis. There were only a few cases when the same plant species grew in both habitats, these cases were excluded from the analysis.

### Plant species importance based on simulations of coextinctions

The effect of extinctions of individual plant species on coextinctions of pollinators was simulated for each of the seven networks with pooled observations from the sown flower strip and the semi-natural habitat (iii). The aim of the simulations was to find out how the community of wild bees responds to the loss of food resources, to quantify the importance of individual plant species for the robustness of the plant-pollinator community, and to test how the plant importance differs between plants from the sown flower strips and semi-natural habitat and depends on other plant characteristics. We carried out the simulations using the two coextinction models described above, i.e TCM and SCM.

Simulations using the TCM were done using the functions second.extinct() and robustness() in the ‘bipartite’ package for R (Dormann et al. 2008). For each network, we generated 10000 random permutations of the plant order and simulated pollinator coextinctions in response to losing plants in that random order. For each simulation, we calculated the estimate of the robustness (Burgos et al. 2007) using the robustness() function of the ‘bipartite’ package. We followed the approach of Pocock et al. (2012) to calculate the contribution of individual plant species to the network robustness, i.e. the plant importance index. For each plant species, we calculated the correlation between the order of its extinction and the network robustness in individual simulations. The coefficient of determination is used as the plant importance index. The justification is that when plants which are important for the network robustness go extinct later in the extinction sequence, multiple pollinator coextinctions are triggered, see Pocock et al. (2012) for details.

The simulations under the SCM conditions were done with a script provided by Vieira & Almeida-Neto (2015). The model has one free parameter *Ri* for each species *i*, which expresses the level of dependence of the species on its interaction partners in the network. Bees are strongly dependent on floral resources, so we assumed that they would disappear with a high probability after losing the plants they visit. Accordingly, we set high values of *Ri* for pollinators, sampled randomly from the interval 0.6 ≤ *Ri* ≤ 0.9 as suggested by Vieira & Almeida-Neto (2015). On the contrary, we assumed that plant species can survive for some time even without pollinators. Hence, we set intermediate values of *Ri* for plants, sampled randomly from the interval 0.3 ≤ *Ri* ≤ 0.6 following Vieira & Almeida-Neto (2015). We run 10000 simulations of plant extinctions in a random order, recorded the order of extinctions, and calculated the network robustness using the same method as for the TCM (Burgos et al. 2007). We then also calculated the plant importance index the same way (see above).

We used generalized linear mixed effect models (GLMM) with Gaussian error distribution and site identity as a factor with random effect to compare the plant importance values between plants unique to sown flower strips and plants unique to semi-natural habitats as for other species-level measures. We also tested whether the values of the plant importance index depended on the three species-level metrics calculated based on network structure, i.e. ND, d’, and PSI (see above).

## Results

In total, we observed 3121 interactions between 47 species of plants and 42 species of Aculeata of seven families (Andrenidae, Apidae, Colletidae, Crabronidae, Halictidae, Megachilidae, and Melittidae). The most abundant were Apidae, in particular the honeybee, *Apis mellifera*. Supporting Information including raw data (Supporting Information Table S1) is available in Figshare at https://doi.org/10.6084/m9.figshare.14185160.

### Species richness and network structure

Semi-natural habitats at the study sites provided on average 10.7 flowering plant species visited by insects, while sown flower strips were 61% poorer in plant species richness, with the mean of 4.2 plant species (t = -5.437, df = 6, p = 0.002, paired t-test on log-transformed data). Correspondingly, the number of bee species was on average 42% lower in the sown flower strips than in the semi-natural habitats (mean 5.6 and 9.7 species, respectively; t = -4.604, df = 6, p = 0.004, paired t-test on log-transformed data). In addition, sown flower strips were dominated by *Apis mellifera*, which accounted for ca. 80% of all individuals of Aculeata, but had a lower percentage of bumblebees (*Bombus* spp.) and particularly solitary bees compared to semi-natural habitats (Table 2). We include Halictidae, which contain also primitively eusocial species, among ‘solitary bees’ in line with previous studies (see e.g. Wood et al. 2017).

**Table 2.** The average percentage of flower visits per plot by honeybees, bumblebees, and solitary bees in sown flower strips and semi-natural habitats and the range of values of individual sites.

| Habitat type | *Apis mellifera* | *Bombus* spp. | Solitary bees |
| --- | --- | --- | --- |
| Sown flower strips | 76.3% (43.3-97.4%) | 17.3% (2.0-35.2%) | 6.4% (0-21.5%) |
| Semi-natural habitats | 55.0% (24.3-87.8%) | 25.3% (9.6-63.3%) | 19.1% (2.2-42.4%) |

Bees also had higher beta-diversity across the seven sites in semi-natural habitats than in sown flower strips. The Shannon index of bee beta-diversity was 2.556 in semi-natural habitats and 1.837 in sown flower strips. This difference apparently follows from the higher beta-diversity of plants across sites in the semi-natural habitats compared to the sown flower strips which were initiated by the same seed mixture. Shannon index of plant beta-diversity was 3.219 for semi-natural habitats compared to 2.383 for sown flower strips.

Differences in the network structure between the sown flower strips and semi-natural habitats were modest, but networks in the sown flower strips were more generalised and nested (Fig. 2, Supporting Information Table S2. We did not find any difference in the level of web asymmetry, i.e. the relative difference between bee species richness and plant species richness, between the sown flower strips and semi-natural habitats (t = 2.167, df = 6, p = 0.073, paired t-test). However, nestedness estimated as weighted NODF was higher in the sown flower strips than in the semi-natural habitats (mean of the differences = 0.74, t = 4.72, df = 6, p = 0.003, paired t-test on log-transformed data). On average, the wNODF was 15.835 in sown flower strips and 33.155 in semi-natural habitats. The community-level degree of specialization (H’2) was also higher in the semi-natural habitats than in the sown flower strips (mean of the differences = 0.244, t = -3.044, df = 6, p = 0.023, paired t-test) with the average value of 0.247 in sown flower strips and 0.492 in semi-natural habitats.

**Fig. 2.**
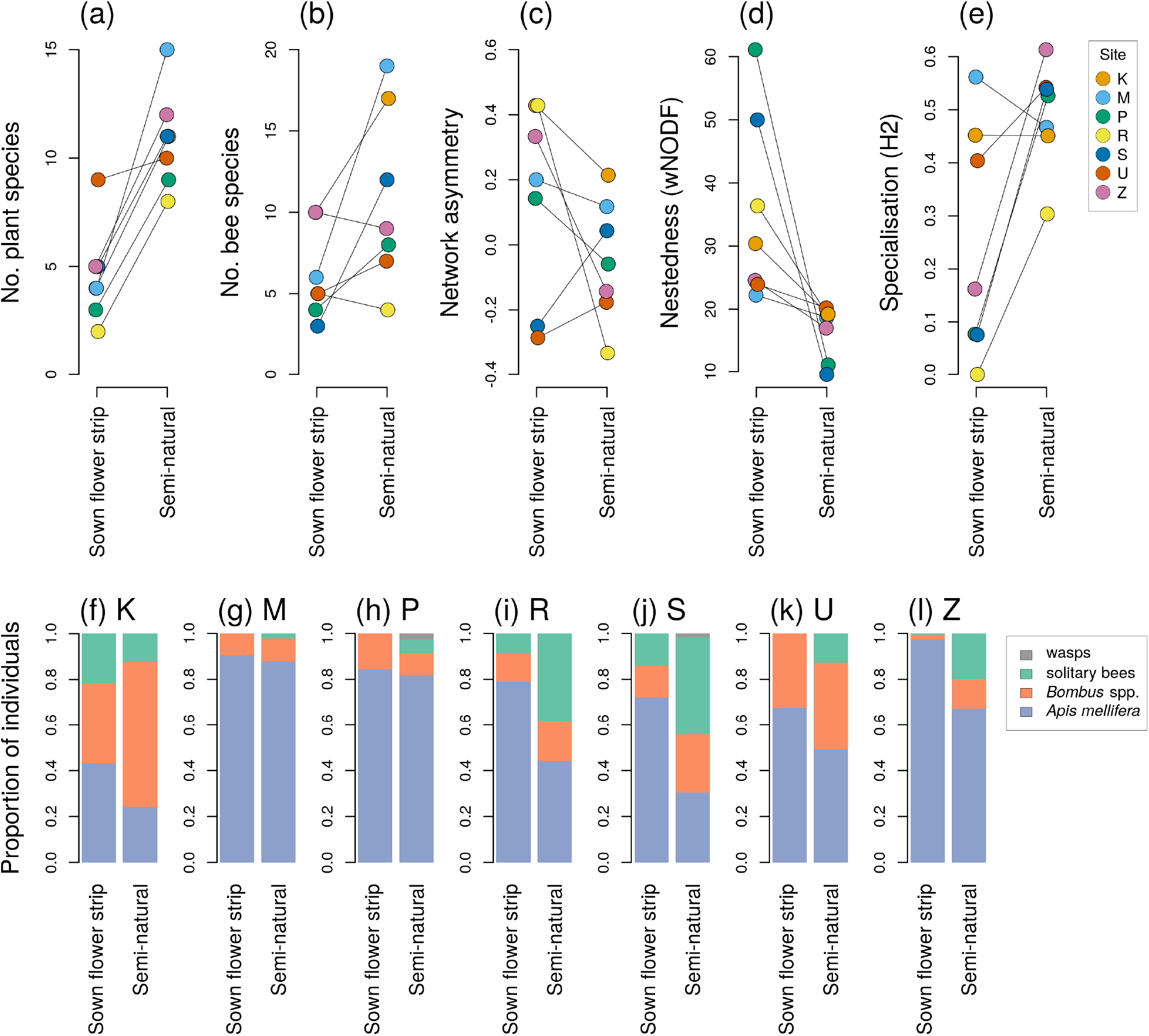
Metrics of plant-pollinator network structure (a - e) and taxonomic composition of pollinators (f -l) in the sown flower strips and semi-natural habitats in individuals sites. See the Methods for details on the network metrics.

However, the robustness (R) of the networks to the secondary extinctions, measured as the area under the coextinction curve, under the conditions of both coextinction models (TCM and SCM) differed very little between sown flower strips and semi-natural habitats (Fig. 2, Supporting Information Table S2). Robustness to simulated extinctions of plants with secondary extinctions of pollinators was similar in both habitat types based on the TCM (mean of the differences = 0.011, t = 0.563, df = 6, p = 0.596) as well as SCM (mean of the differences = -0.008, t = -2.105, df = 6, p-value = 0.08). Robustness to simulated extinctions of pollinators with secondary extinctions of plants also did not differ between the two habitats according to TCM (mean of the differences = 0.03, t = 1.126, df = 6, p = 0.303). However, there appeared to be a significant difference in robustness to simulated extinctions of pollinators according to SCM, with slightly higher robustness in the semi-natural habitat, but the difference was very small and biologically negligible (mean of the differences = -0.014, t = -2.486, df = 6, p = 0.047).

### The importance of plants in the sown flower strips and semi-natural habitats for bee diversity

In general, we found no significant differences between species-level indices of plants from sown flower strips and semi-natural habitats (Table 3). However, there was high interspecific variance in plant species-level indices suggesting that individual plant species vary substantially in their importance for the local pollinator community (Supporting Information Table S3).

**Table 3.** A comparison of the values of species-level indices of plants associated with the sown flower strips and semi-natural habitats. Species-level indices were calculated using pooled networks combining data from both habitats in individual sites. Plants species were classified as occurring only in the sown flower strips or only in the semi-natural habitats. Mean values and standard errors (SE) are shown for both groups of plants, together with the results of generalised linear mixed models (GLMM) comparing the values of species-level indices between plants associated with the sown flowering strips and semi-natural habitats.

| Species-level index | Sown flower strips | | Semi-natural habitats | | $\chi^2$ | P |
| --- | --- | --- | --- | --- | --- | --- |
|  | mean | SE | mean | SE |  |  |
| Log(normalised degree) | -1.74 | 0.196 | -1.88 | 0.162 | 0.72 | 0.3968 |
| d' | 0.16 | 0.058 | 0.28 | 0.034 | 3.07 | 0.0796 |
| Log(PSI) | -2.71 | 0.378 | -2.81 | 0.225 | 0.05 | 0.8213 |
| Log(Plant importance – TCM) | -4.40 | 0.440 | -3.89 | 0.274 | 1.08 | 0.2980 |
| Log(Plant importance – SCM) | -4.26 | 0.390 | -4.57 | 0.235 | 0.45 | 0.5045 |

The plant importance index based on the effect of simulated plant extinctions varied among species, but did not consistently differ between plants of sown flower strips and semi-natural habitats (Fig. 3, Supporting Information Table S3). The results were similar for both coextinction models (TCM and SCM). There was only a weak relationship between the plant importance index and other species-level indices (Table 4). The plant importance index based on the SCM was higher in plant species which had higher relative visitation rate, but only marginally higher in plants with a higher normalised degree, i.e. relative number of flower visitor species. These trends were not statistically supported for the plant importance index based on coextinction simulations by TCM (Table 4).

**Fig. 3.**
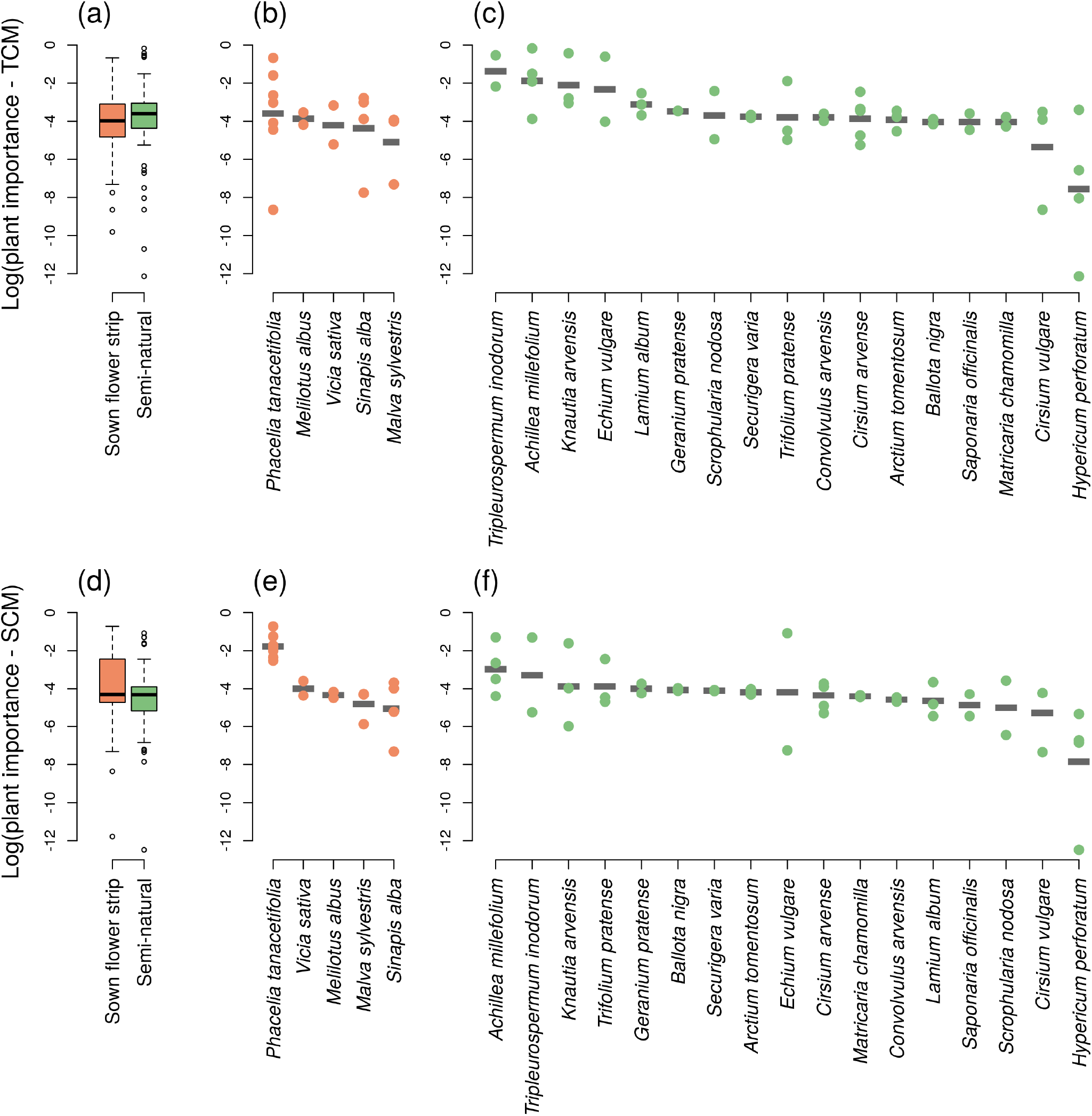
A comparison of values of the plant importance index based on simulations of coextinctions using two coextincion models, TCM and SCM. Plant importance values across all plant species in the two habitats are shown based on the TCM (a) and SCM (d), along with plant importance values of individual plant species exclusive to the sown flower strips (b and e) and semi-natural grasslands (c and f). Only plant species present at two or more sites are shown in b), c), e), and f).

**Table 4.** The dependence of the plant importance index in the two habitats on the relative visitation, normalised degree, and specialisation (d’) of individual plant species.

| Predictor | Plant importance (TCM) |  |  |  | Plant importance (SCM) |  |  |  |
| --- | --- | --- | --- | --- | --- | --- | --- | --- |
| | Estimate | SE | $\chi^2$ | P | Estimate | SE | $\chi^2$ | P |
| Log(relative visitation) | 0.17 | 0.124 | 2.05 | 0.1524 | **0.34** | **0.109** | **9.30** | **0.0023** |
| Log(normalised degree) | 0.48 | 0.291 | 2.69 | 0.1009 | 0.51 | 0.263 | 3.30 | 0.0694 |
| d' | -0.35 | 0.722 | 0.20 | 0.6536 | -0.81 | 0.653 | 1.52 | 0.2173 |

Among the sown plant species, *Phacelia tanacetifolia* was the most visited plant in sown flower strips at all sites. This species was also overall the most visited plant species at all sites except one, where naturally occurring *Echium vulgare* had even more visits. *Phacelia tanacetifolia* had also relatively high values of species-level indices based on the structure of the plant-pollinator networks (PSI and ND), low degree of specialisation (d’), and high level of the plant importance index based on simulations of coextinctions using both coexinction models (Fig. 3, Supporting Information Table S3). However, its primary visitors were large generalist social bees such as the honeybee (*Apis mellifera*) and bumblebees (*Bombus* spp.), while it was less visited by solitary bees (Supporting Information Table S1).

## Discussion

A considerable effort has been directed towards creating suitable habitats for animals, particularly pollinators, in agricultural landscapes over the past two decades, often within the administrative framework of Agri-Environment Schemes (AES) (Haaland et al. 2011), and later Agri-Environment Climate Schemes (AECS). Sown flower strips are only one of the measures introduced, but they have become particularly promoted. However, our results show that plant-pollinator interaction networks in the sown flower strips had lower alpha as well as beta diversity of plants and pollinators in comparison with networks from semi-natural habitats in their surroundings. On the other hand, sown flower strips contained nectar-rich and thus highly visited plant species, mainly non-native *Phacelia tanacetifolia*, which resulted in higher level of nestedness and lower level of specialisation of plant-pollinator interactions in sown flower strips compared to semi-natural habitats. Overall, the sown flower strips supported high abundance of aculeate Hymenoptera, but the vast majority were common generalist species, such as honeybees which made up almost 80% of individuals in the sown flower strips. More specialized solitary bees, and to a lesser degree also bumblebees, were more abundant and had higher species richness in nearby semi-natural grasslands.

Unlike most previous studies, which compared the abundance and diversity of pollinators (or other invertebrates) between sown flower strips and agricultural crops (usually wind-pollinated, such as cereals), our study directly compared sown flower strips with semi-natural habitats in the landscape matrix among the fields. It is well established that sown flower strips host a higher abundance and diversity of pollinators compared to most crops because they offer a more abundant and diverse floral resources compared to e.g. cereal fields (reviewed by Haaland et al. 2011). Creating sown flower strips may thus benefit pollinators in areas with very intensive agriculture (Buhk et al. 2018), but may be of limited value in more extensive landscapes with sufficient cover of semi-natural habitats (Tscharntke et al. 2011, Scheper et al. 2013). Our results support this suggestion. While out sites were located in productive agricultural landscapes, there were still extensive areas of semi-natural habitats among the fields (Fig. 1), and only a minority of aculeate Hymenoptera, particularly solitary bees, occurring there were regularly found in the sown flower strips. In accordance with our findings, Wood et al. (2017) observed that only a minority of solitary bee species found at multiple farms in southern England visited sown plant species and collected their pollen, while most solitary bee species were restricted to wild plants.

Even though the size and mowing regime of sown flower strips at different sites varied, they all hosted similar communities of bees with lower beta-diversity compared to the semi-natural habitats. This may be partly caused by identical seed mix used for sown flower strips at all sites. However, the beta-diversity of pollinators may be affected also be affected by phenology (Carvell et al. 2006, Ouvrard et al, 2018) and heterogeneity of vegetation structure because pollinators tend to modify their preferences for flowers based on the characteristics of vegetation such as the composition of surrounding flowers (Janovský et al. 2013) or the height of the surrounding vegetation (Klečka et al. 2018a). Despite variable mowing regimes, such structural patterns of the vegetation were still more variable in semi-natural habitats than in sown flower strips.

Our application of approaches of ecological network analysis provided novel insights into the importance of sown and wild plants for conservation of wild bees in agricultural landscapes. We found considerable differences in the structure of plant-pollinator interaction networks from sown flower strips compared to semi-natural habitats. In particular, the communities from sown flower strips tend to have higher level of nestedness and lower level of specialisation, altogether indicating less complex communities built around only a few central species such as *Phacelia tanacetifolia* and *Apis mellifera*. However, both types of communities were very similar in their robustness to secondary extinctions triggered by primary extinctions of plants as well as pollinators, simulated by two different types of models (TCM and SCM). Yet, the communities still differ in the level of specialisation of plant-pollinator interactions which may affect their reaction to disturbances (Vázquez & Simberloff 2002).

In the history of theoretical ecology, a relationship between complexity of a community and its stability has been challenging for a long time (Pimm et al. 1984). Nowadays, using the analyses of interaction networks, terms of complexity and stability can be accessed in much more details. Here, we show that even when the complexity of communities (i.e. diversity of species as well as entropy of their linkage H’2) was significantly higher in semi-natural habitats than in sown flower strips, their stability (i.e. robustness to secondary extinctions) did not differ much between the two habitats. However, the applicability of the analyses of community stability or robustness for the purpose of conservation biology remains an open question (Tylianakis et al. 2010).

Enhancing the bee diversity in flower strips in the year of sowing (the case of our study) may be possible by enriching the seed mixture, especially if annual plant species from the Asteraceae family are included (Nowakowski & Pywell 2016, Ouvrard et al. 2018). Asteraceae species may be responsible for the higher bee beta-diversity in semi-natural habitats compared to flower strips and were represented mainly by *Tripleurospermum inodorum, Tanacetum vulgare, Achillea millefollium, Centaurea* spp., *Cirsium* spp., alongside with Lamiaceae species (e.g. *Ballota nigra, Lamium album, Lamium purpureum*), *Echium vulgare* (Boraginaceae), *Knautia arvensis* (Caprifoliaceae), and more species belonging to other botanical families. Increasing the floral diversity of the sown flower strips would likely lead to an increase of the diversity of solitary bees (Wood et al. 2017) and other pollinators. Most pollinators display preferences for various floral traits, such as flower size, colour, and shape (Junker et al. 2013, Klečka et al. 2018b), or vegetation structure, such as plant height (Klečka et al. 2018a) and spatial clustering (Janovský et al. 2013, Blaauw & Isaacs 2014, Akter et al. 2017). Increasing the functional diversity of plants in the sown flower strips and their microhabitat heterogeneity might have beneficial effects on pollinator communities by providing suitable conditions for a broader range of species and by decreasing their competition with dominant species such as *Apis mellifera* (Forup & Memmott 2005, Hudewenz & Klein 2013). Moreover, some species recorded in sown flower strips may visit flowers of sown plants only as a source of nectar, but require additional plants as pollen source necessary for feeding their larvae (Wood et al. 2017). Furthermore, higher diversity of plants would ensure the continuity of the presence of sources of nectar and pollen from early spring until the end of summer. Such temporal continuity of floral resources has been recognised as a key factor for the persistence of a species rich community of pollinators (Scheper et al. 2015, Wood et al. 2017, Gayer et al. in press).

While our study focused on bees, other groups of insects such as Diptera are important pollinators too (Larson et al. 2001, Ssymank et al. 2008, Klečka et al. 2018b). The species composition of seeds used for sown flower strips in our study was designed specifically to support bees and plant species were chosen mainly to provide high amount of nectar. Plants offering high amount of nectar may be favoured not only by bees, but also by butterflies and some groups of long-tongued Diptera (Labandeira 2000). On the other hand, other groups of flower visiting insects such as Coleoptera and many groups of Diptera or parasitic Hymenoptera visit flowers mainly for pollen (Labandeira 2000), so different flowering species such as yellow Asteraceae and Apiaceae may be more important food source for these groups of pollinators (Hegland & Boeke 2006, Honěk et al. 2016, Klečka et al. 2018b). Although different groups of pollinators may vary in their floral and habitat requirement, we suggest that the exclusive focus of our study on aculeate Hymenoptera is justified for several reasons: i) sown flower strips were designed mainly to support bees; ii) in comparison with many other groups of pollinators, bees usually have a relativelly narrow diet and may thus be relatively sensitive to inappropriate selection of sown plants; iii) in bees, not only adults, but also larvae are reliant on food from flowers, so the pollination network should also reflect the food web of bee larvae; and iv) aculeate Hymenoptera is a diverse group with a number of species of conservation relevance with specialised requirements of floral resources, climatic and habitat conditions, and high degree of philopatry because of their nesting strategy, so they can serve as proper bioindicators at small spatial scales (Tscharntke et al. 1998, Talašová et al. 2018).

Flower-rich vegetation is favourable not only for pollinators, but also other economically important groups of insects such as natural enemies of pests (Brennan 2013, Tschumi et al. 2016, Karamaouna et al. 2019). Designing flower strips in such a way to satisfy requirements of multiple groups of animals and to provide multiple ecosystem services beyond conservation of insect diversity requires a careful selection of plant species based on multiple criteria (Cresswell et al. 2019). The approach to quantifying plant species importance based on simulations of coextinctions in plant-pollinator networks, as used in our study, can be readily extended to other groups of animals interacting with the plants (Pocock et al. 2012). Importantly, Pocock et al. (2012) found that the importance of plant species for insect pollinators is not a good predictor of its importance for different insect guilds, so further research will be needed to test the suitability of sown flower strips for other taxonomic and ecological groups of insects. However, there is already evidence that open flower plants such as Asteraceae or Apiaceae are particularly appropriate for the purpose of hosting highly diverse community of animals from many different taxa and trophic groups (Pocock et al. 2012). In our sown flower strips, only two species from these families were sown: *Achillea millefolium* (Asteraceae) and *Carum carvi* (Apiaceae), and no bee pollinator was recorded on them, perhaps due to low densities of these plants during the first year after the sowing.

Our results confirm that sown flower strips may be a useful conservation measure, but we highlight that appropriate management of semi-natural habitats maximising their floristic and functional diversity is still indispensable. This includes diversified mowing regimes of meadows (Čížek et al. 2012), management of transitional habitats at the borders of meadows and forests (Slámová et al. 2013), and low-intensity grazing (Kruess & Tscharntke 2002, Sjödin et al. 2008, Zhu et al. 2012). As we have shown, semi-natural habitats may support a broader range of pollinator species. However, sown flower strips with a functionally diverse selection of plant species may still represent a valuable complementary conservation measure in areas with intensive agriculture (Tscharntke et al. 2011, Scheper et al. 2013, Buhk et al. 2018).

## Conclusions

Sown flower strips have become a popular tool for supporting the communities of pollinators such as aculeate Hymenoptera. Our results show that while they may support high bee abundances, they are visited by a restricted set of species, mostly abundant generalists, compared to nearby semi-natural habitats. Hence, sown flower strips themselves are not sufficient for the preservation of the diversity of pollinating insects. We suggest that enhancing flowering plant diversity and enhancing the diversity of structural properties of the sown flower strips could increase their potential for maintenance of the biodiversity of pollinators. Our novel analysis of plant species importance based on simulations of coextinctions in plant-pollinator networks shows large variance of plant importance scores among species, which suggests that strategic selection of plant species may increase the conservation values of sown flower strips. Nevertheless, conservation of natural and semi-natural habitats with diverse and heterogeneous vegetation will still play a key role in the conservation of bee diversity and their appropriate management should be given priority, particularly in landscapes where such habitats remain common.

## Acknowledgements

The work of JH was supported by the Institutional Research Support grant of the Charles University, Prague (No. SVV 260 434/2018). The study was also supported by the Czech Science Foundation (projects GJ17-24795Y and GA20-14872S). We thank Tomáš Frýda for his comments on statistical analyses.

## Data availability statement

The data that support the findings of this study are openly available in Figshare at https://doi.org/10.6084/m9.figshare.14185160.

## Supporting information

Supporting Information (Tables S1-S3) is available in Figshare at https://doi.org/10.6084/m9.figshare.14185160.

## Notes

https://doi.org/10.6084/m9.figshare.14185160

### Competing Interest Statement

The authors have declared no competing interest.

